# Sex-dependent Lupus *Ruminococcus blautia gnavus* strain induction of zonulin-mediated intestinal permeability and autoimmunity

**DOI:** 10.1101/2021.07.06.451365

**Authors:** Jing Deng, Doua F. Azzouz, Nicole Ferstler, Gregg J. Silverman

## Abstract

Imbalances in the gut microbiome are suspected as contributors to the pathogenesis of Systemic Lupus Erythematosus, and our studies and others have documented that patients with active Lupus nephritis have expansions of the obligate anaerobe, *Ruminococcus blautia gnavus* (RG). To investigate whether the RG strains in Lupus patients have *in vivo* pathogenic properties, we colonized C57BL/6 mice with individual RG strains from healthy adults or those from Lupus patients. These strains had a similar capacity for murine intestinal colonization, in antibiotic-preconditioned specific-pathogen-free, as well as germ-free adults, and their neonatally colonized litters. Lupus-derived RG strains induced high levels of intestinal permeability that was significantly greater in female than male mice, whereas the RG species-type strain (ATCC29149/VPI C7-1) from a healthy donor had little or no effects. Lupus RG strain-induced functional alterations were associated dysregulated occluden transcript production in the ileal wall as well as raised serum levels of zonulin, a regulator of tight junction formation between cells that form the gut barrier. Notably, the level of Lupus RG-induced intestinal permeability was significantly correlated with serum IgG anti RG cell-wall lipoglycan antibodies, and to anti-native DNA autoantibodies that are a biomarker for SLE. Strikingly, gut permeability was completely reversed by oral treatment with larazotide acetate, an octapeptide that is a specific molecular antagonist of zonulin. Taken together, these studies document a molecular pathway by which RG strains from Lupus patients induce a leaky gut and autoimmunity that have been implicated in the pathogenesis of flares of clinical Lupus disease.

## Introduction

Our gut microbiomes contain a complex interdependent community that in health provides layered tiers of nutritional and immune regulatory benefits. Imbalances (or dysbiosis) have been implicated in a growing number of clinical conditions but only in a handful of cases have expansions of specific bacteria been implicated, and in even fewer cases have actual pathogenic pathways been identified. In studies of a clinically diverse cohort of female patients with Systemic Lupus Erythematosus (SLE), we discovered the first correlation between overrepresentation of an obligate anaerobe, *Ruminococcus blautia gnavus* (RG) with SLE disease activity (1). More intriguingly, even amongst Lupus patients these RG expansions were more common in patients with Lupus nephritis, which effects more than half of patients and is associated with amongst the greatest morbidity and mortality (1). Notably, the association of RG with LN was also later independently documented in a large cohort of untreated Chinese Lupus patients (2). In more recent longitudinal studies of our cohort, Lupus patients were found to have inherently unstable gut microbiota communities (3) and almost half of clinical flares of renal disease were temporally associated with ephemeral RG blooms (3).

One of the estimated 53 most common human intestinal colonizers, *RG* are early colonizers that are detectable in most infants by 24 months of age (4). In adults, RG is present in at least 90% of individuals from North American and Europe, although generally at stable low-levels at or below 1% abundance (5, 6). Based on genomic phylogenetic analysis, RG has been reassigned to the genus Blautia within the Lachnospiraceae family of spore-forming obligate anaerobes. As a species, RG is quite distinct from other taxa at both the genome as well as for 16S rRNA gene sequence level (7). In healthy adults, RG is a keystone species present in 90% or more of adults (5, 6), playing pleotropic roles in host metabolism and immunity (reviewed in (8)), including for the conversion of primary to secondary bile acids (9), production of the short-chain fatty acids (SCFAs) that aid immune regulation (10).

Most reports of the disease-associated variations in microbiota communities have been limited to correlative studies, and with limited exception (11), the *in vivo* effects of RG isolates on host immunity have largely focused on the Human Microbiome Project designated type-specific RG strain, VPI C7-9, also termed ATCC29149 (and referred to in our studies as RG1) (12–14) that was isolated from stool of a healthy donor (15). While increased intestinal abundance of this species alone might explain associations with Lupus disease flares, we hypothesize that there may be important differences in the pathogenetic potential of strains from healthy individuals from those colonizing LN patients. We therefore set out to compare the effects of different genome-defined RG strains on the host following *in vivo* intestinal colonization, with an emphasis on whether gut barrier function was affected. Whereas we found that all RG strains we evaluated had the capacity to colonize the mouse gut, there were dramatic differences on their effects on intestinal permeability, which was found to mediated by a zonulin-dependent mechanism. Strains isolated from clinically active Lupus patients reproducibility induced these changes. Moreover, neonatal murine colonization with Lupus RG strains resulted in microbial translocation and systemic antibody responses to RG-specific antigens and induction of Lupus autoantibodies. These findings provide a mechanistic rationale for the previously reported linkage between RG intestinal expansions and Lupus pathogenesis.

## Methods

### Mice

All animal work was performed under the supervision of the NYU Langone IACUC. Germ-free (GF) mice were bred and maintained in the gnotobiotic facility at the NYU Langone (16). All mice were C57BL/6 genotype, and locally bred or purchased from Charles River Laboratories (Wilmington MA) and were received at 6–8 weeks of age, or locally bred. All other mice were, maintained in specific-pathogen-free (SPF) cages, with free access to food and water. To avoid possible cross-contamination mice colonized with each of the different strains were separately raised in isolator cages. Mice were housed under a 12-h light/dark cycle at 23 °C.

### Intestinal colonization with RG strain

Individual *RG* strains were streaked then individual colonies grown in 5 ml of BHI media (Anaerobe Systems) under anaerobic conditions overnight. GF C57BL/6 mice (4 male and 4 female) were colonized by oral gavage of different *RG* strains (RG1, RG2 and S107-48) of 10^8^ CFU in 500 μL of sterile PBS, every other day for a total of five times. Individual mice were weighed, then fecal pellets and bleeds were collected from individual mice prior to gavage, and again at days 7, 14, and 21 following gavage. Pellets were stored at −80 °C until DNA extraction. The bacterial translocation and burden of *R. gnavus* in feces was determined by RG-specific qPCR at the indicated time points. All breeding airs yielded litters that were then housed under SPF conditions.

To colonize mice previously raised under Specific Pathogen-Free (SPF) conditions, 4-6 week old mice were preconditioned with oral antibiotics, then breeding pairs were gavaged with RG1, RG2, or Lupus strains S107-48 or S47-18, and only the pair colonized with the RG1 strain did not yield litter(s). The antibiotic cocktail was composed of vancomycin (0.5g/L)(Fisher Scientific), neomycin (1g/L)(Fisher Scientific), ampicillin (1g/L; Fisher Scientific) and metronidazole (1g/L; Fisher Scientific), with solutions freshly prepared each week in autoclaved drinking water, and all antibiotics remained soluble at this concentration. Antibiotics were provided in 100-ml clear glass sippers (Braintree Scientific, Inc., Braintree MA). SPF mice received systemic antibiotics (see above) at 4 weeks of age for one month. Fecal pellets were collected prior to, and following antibiotic exposure at days 21, and then weekly until at least 100-fold decreasing of total 16S rRNA DNA (representing bacterial burden) in fecal pellets by qPCR was confirmed from each mouse. Mice were then switched to plain water *ad libitum* for 24 hours, then mice then received the same as above described oral gavage of different *RG* strains (RG1, RG2, S107-48 and S47-18)(see Supplementary Figure 1).

### Quantitative PCR analysis

For bacterial genomic 16S rRNA gene quantitation, DNA was isolated from mice fecal pellets, cecum contents, spleens and mesenteric lymph nodes (MLNs) using QIAamp DNeasy Powersoil or Blood & Tissue kit (Qiagen), according to the manufacturer’s instructions. DNA isolated from all these sample was quantified on a Nanodrop 1000 (Thermo Scientific) and then run for quantitative PCR assays on the StepOnePlus™ Real-Time PCR System (Applied Biosystems) with the Power SYBR Green Master Mix (Applied Biosystems). For end-point PCR reactions, a thermal cycler (Applied Biosystems) was programmed to amplify bacterial DNA using the following primers that were also used for qPCR experiments. PCR reactions were run with the following conditions: holding stage at 95°C for 5 minutes followed by 50 cycles of 95 °C for 15 seconds, 58 °C for 30 seconds; and followed by extension at 72°C for 5 minutes. Total bacterial 16S rRNA gene content was assessed with the oligonucleotide primers:

UniF340(5’-ACTCCTACGGGAGGCAGCAGT-3’)
UniR514(5’-ATTACCGCGGCTGCTGGC-3’).

The RG species-specific 16S rRNA was determined with previously reported oligonucleotide primers (17):

Fwd 5’-GGACTGCATTTGGAACTGTCAG-3’
Rev 5’-AACGTCAGTCATCGTCCAGAAAG-3’

For host gene transcript quantitation, RNA was isolated from ileum using the RNAeasy plus mini kit (Qiagen) according to the manufacturer’s instructions. The RT-cDNA was generated using the Invitrogen™ SuperScript™ II Reverse Transcriptase (ThermoFisher), according to the manufacturer’s protocol. RT-qPCR was performed on cDNA that was amplified using gene specific primers, with Power SYBR Green Master Mix (Applied Biosystems), and the StepOnePlus™ Real-Time PCR System (Applied Biosystems). After normalization to GAPDH levels, each sample gene expression was quantified as the -fold increase from baseline sample. All real-time PCR reactions were performed in triplicate. For RT-qPCR, the following murine-specific oligonucleotide primer sequences were used:

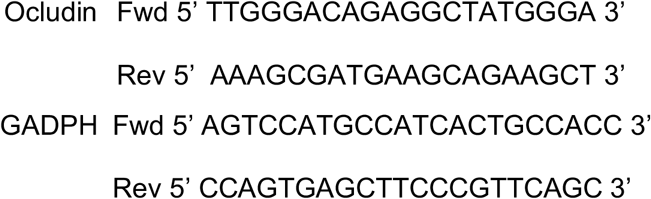

### In vivo assay of intestinal permeability

To assess intestinal permeability, after 4 h fasting of food and water, mice were orally gavaged with 4,000-Da fluorescein (FITC)-dextran (FD4) (Sigma-Aldrich, St. Louis, MO) (250 mg/kg body weight) in doses of 200 μl, and blood was collected 3 hr later. The concentration of the FD4 was determined using a fluorimeter with an excitation wavelength at 490 nm and an emission wavelength of 530 nm. To assess concentration, FD4 in serum was then serially diluted to establish a standard curve.

#### Leaky gut recovery study

Zonulin antagonist, larazotide acetate (known as AT-1001, or INN-202), was purchased from BOC Sciences, NY). Individual neonatally-colonized littermates from SPF mice, colonized with different RG strains, were retested after FD4 challenge, and those with abnormal levels each then received 0.15 mg/ml of the zonulin antagonist in the drinking water, which was refreshed every day, for 10 consecutive days. After 24 hr rest, intestinal permeability was then retested.

### Assays of antigen-reactive IgG antibodies

A custom multiplex bead-based array for the Magpix platform (Luminex, Austin TX) was created by coupling a variety of highly purified thymic native DNA, purified cell wall lipoglycan (LG) from the RG2 strain (i.e., IgG anti-RG2 LG (formerly termed LG3 (1)), and the Lupus RG strain S47-18 (i.e., IgG anti-S47-18 strain LG), recombinant *S. aureus* proteins, endotoxins, and other bacterial antigens and control ligands, to individual microspheres, adapting the manufacturer’s protocol and previously reports (18–20). For antigen-reactive IgG detection, 1,000 microspheres per analyte per well were premixed, sonicated, and then with addition of 100 μl of serum diluted, as indicated. For ELISA bound IgG antibodies were detected with Fc-gamma-specific anti-murine IgG HRP (eBioscience, San Diego CA), and for multiplex bead-based assay with anti-mouse IgG (Fc-specific) *F*(ab’)2 PE (Jackson ImmunoResearch, West Grove PA). Data were acquired on a Magpix instrument (Luminex) and reported as mean fluorescence intensity (MFI) values, as previously described (19–21). Plasma zonulin were quantitated with a commercial ELISA Kit. (MyBioSource, San Diego CA.)

### Biostatistics

Comparisons were with two-tailed unpaired or paired t-tests, or Spearman correlations, as indicated, with Prism version 9.0 for Mac OS (Graphpad, San Diego CA). p<0.05 was significant.

## Results

### RG persistence in germ-free mice with transmission to litters

In these studies, we sought to investigate for differences in the *in vivo* pathogenicity properties of *Ruminococcus blautia gnavus* (RG) strains isolated from a fecal sample of a healthy donor, VPI C7-9 (here termed RG1) and from a colonic biopsy, CC55_001C (here termed RG2), with comparisons to the RG strains, S107-48 and S47-18 obtained from fecal samples of two SLE patients with RG blooms at the time of flare of renal disease (manuscript in preparation). To initiate these studies, we used a standard gavage protocol to colonize groups of germ-free C57BL/6 mice with different RG strains.

Due in part to the practical challenges of accurate CFU determinations from this obligate anaerobe we adapted a previous described RG rRNA species genomic-specific qPCR assay that was validated to be highly RG species specific (17). Whereas in the fecal pellets of naïve germ-free C57BL/6 mice RG was undetectable, following gavage with different RG strains, high levels were detectable at high levels in sequential samples obtained over the four weeks following gavage (Figure 1A). There initially were no differences in levels of fecal RG representation with the different strains, nor differences based on the sex of the recipient mice (Figure 1A and data not shown).

**Figure 1.**
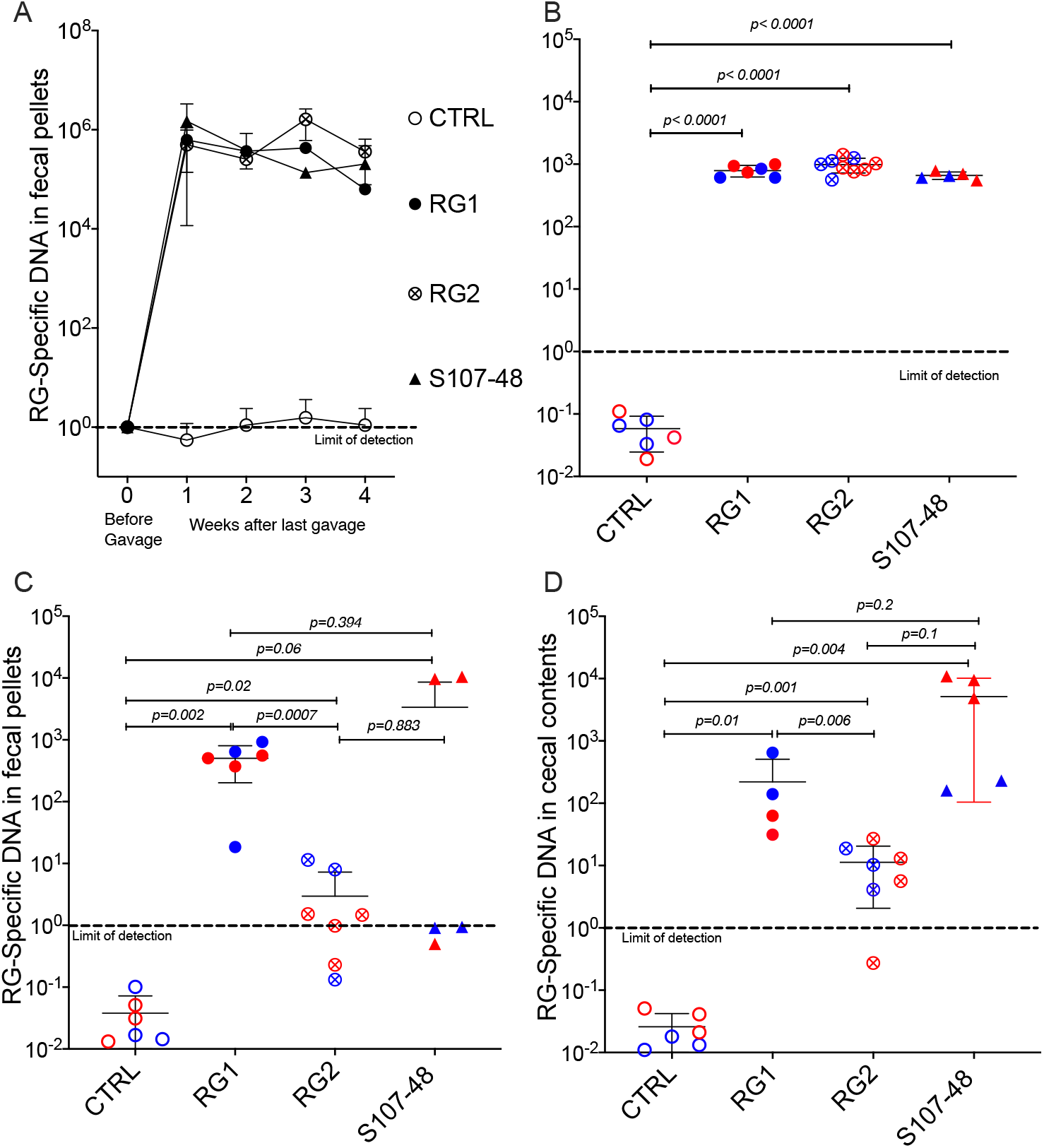
Diverse RG strains isolated from human donors colonize germ-free (GF) mice, and are passed to their litters. A) Abundance of RG DNA in germ-free (GF) C57BL/6 mice breeding pairs, monocolonized by oral gavage with in RG1, RG2 and S107-48 RG strains. RG species-specific 16S rRNA genomic levels were quantified by genomic qPCR analysis of fecal pellets, as shown. Inset values indicate normalized quantity of RG 16S rDNA with control mice, before (left) and weeks after (right) gavage with each of the RG stains, as indicated. B) Abundance of RG-specific genomic DNA in 4wk-old litters of the RG1, RG2 and S107-48-monocolonized GF breeding pairs. Mice. RG species-specific 16S rRNA levels were quantified by genomic qPCR analysis of fecal pellets, as shown. C) Abundance of RG DNA in fecal pellets of 12-14wks-old litters of the individual RG1, RG2 and S107-48 strains monocolonized GF breeding pairs. D) Abundance of RG DNA in cecal pellets of 12-14 wk-old litters of RG1, RG2 and S107-48-monocolonized GF breeding pairs. Note that overtime there is increased variation in RG strain persistence as measured in fecal pellets, but less heterogeneity in RG representation in the luminal extracts of the cecums of these same individual colonized mice. (Red) indicates female and (Blue) male mice are indicated separately.

Earlier studies in mice have demonstrated that adult germ-free mice have immune defects, whereas colonization of neonatal mice results in exposure to antigens of colonizing bacterial species can affect both the host innate immunity, as well as lymphocyte development (22). We therefore mated breeding pairs of these formerly GF mice that had each been colonized with individual RG strains. Each of these strain-colonized pairs yielded litters, which were then weaned, then individual fecal samples were collected and testing showed roughly equivalent high levels of RG-species specific 16S rRNA genes (Figure 1B).

To investigate for the persistence of RG colonization, fecal pellets were also obtained at about 3-months of age, and RG levels were found to diverge based on the colonizing RG strains. In specific, compared to controls highest mean levels were found in mice colonized with the RG1 strain (*p*=0.002), with lower but still significant levels were detected in mice colonized with the RG2 strain (*p*=0.02)(Figure 1C). Strikingly, although each was above the level found in the non-colonized control mice, the RG levels detected in S107-48 Lupus strain colonized mice differed greatly (Figure 1C), and as a consequence did not attain significance compared to controls (p=0.06). Interestingly, in this group fecal RG levels were more heterogeneous, and higher in two of the three females and lower in the two S107-48 colonized male mice (Figure 1C).

In health, Lachnospiraceae are known to differentially colonize regional sites within the intestine (23). Therefore, after sacrifice, the cecal luminal contents were sampled and significant levels of RG-specific genomic DNA were documented in mice colonized with each of the three RG strains (Figure 1D). Notably, the highest mean levels were documented for the Lupus S107-48 strain, followed by the RG1 and then the RG2 strain that had been isolated from healthy individuals. While it was not entirely unexpected that human RG isolates might not persist indefinitely in the murine intestine, it was intriguing to find mice with much higher levels due to cecal persistence at a time of waning RG representation in fecal pellets in many of these same individual mice.

### RG bacteria efficiently colonizes antibiotic-preconditioned SPF mice

We also investigated the capacity of these RG strains to colonize immunocompetent adult mice that had been bred and raised under SPF conditions. To reduce the intestinal bacterial burden, we employed a previously validated combination oral antibiotic regimen and to monitor bacterial depletion we utilized a total 16S rRNA qPCR assay proven to be broadly amplify bacterial taxa of broad phylogenetic origins (see methods) (Figure 2A), By this approach we documented that after a 1 month treatment duration community abundance was reduced at least 100-fold in each of the antibiotic-treated mice (Figure 2A), and then individual mating pairs were then gavaged with different RG strain cultures. In comparisons of paired fecal pellet samples obtained before and two weeks after gavage, although there was inter-individual variation, the RG-specific qPCR assay demonstrated significant levels of RG colonization in each of the recipient mice (Figure 2B). These studies document the capacity of RG to colonize mice raised under SPF conditions.

**Figure 2.**
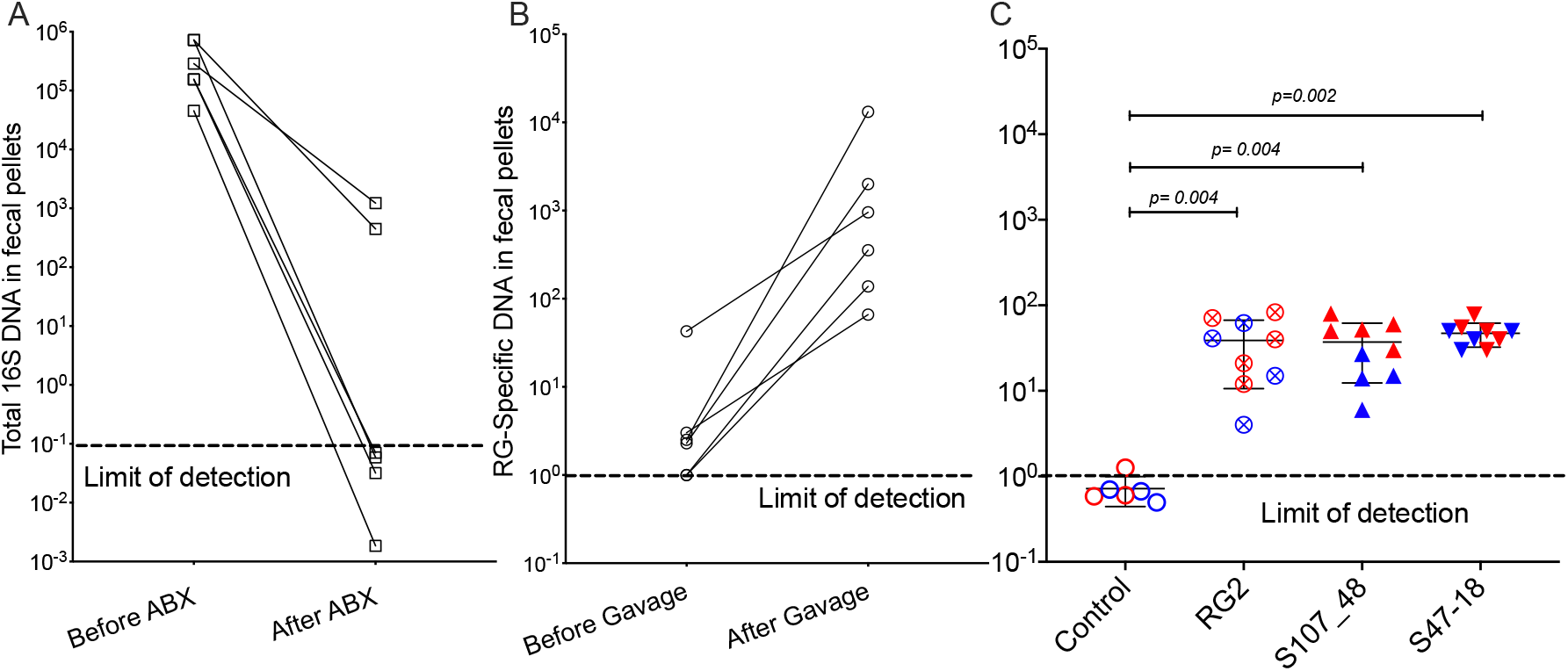
Broad-spectrum oral antibiotic-conditioned mice raised under specific pathogen-free (SPF) conditions are colonized with diverse RG strains isolated from healthy and Lupus-affected human donors. The colonizing RG strains are then passed to their litters. A) The fecal pellets of SPF-raised C57BL/6 breeding pairs were collected before initiating oral regimen of broad-spectrum antibiotics and repeated four weeks later, with total 16S rRNA genomic levels measured by qPCR analysis of extracts of fecal pellets. B) Fecal pellets from these same mice were evaluated for levels of RG-specific genomic DNA, before and after gavage with freshly cultured RG1, RG2 and S107-48 RG strains. C) After mating of pairs based on same RG strain, abundance of RG genomic DNA in fecal pellets of 5wk-old litters was measured in each of the RG1, RG2, S107-48 and S47-18 strain monocolonized SPF littermates, by RG-species specific 16S rDNA genomic DNA by qPCR analysis. (Red) indicates female and (Blue) male mice are separately shown.

### Lupus RG strains induce increased gut permeability

To investigate the potential influences of intestinal colonization on gut barrier integrity, we gavaged individual mice with a standardized dose of FITC-labeled dextran of 4000 Da molecular weight (FD4), then blood samples were subsequently obtained to assess for altered intestinal permeability (Figure 3). In the litters from GF mice colonized with RG, little or no leakage was detected in the weaned pups colonized by RG1. In contrast, there was evidence of increased permeability in both the litters colonized with the RG2 strain and the Lupus strain, S107-48 (Figure 3A). There were no differences in plasma FD4 levels found in male and female mice colonized with the RG1 strain (Figure 3B). Strikingly, the female mice colonized with the RG2 strain and those with the Lupus S107-48 strain had substantially higher levels of post-challenge plasma FD4, consistent with greater intestinal permeability in these RG strains (Figure 3B).

**Figure 3.**
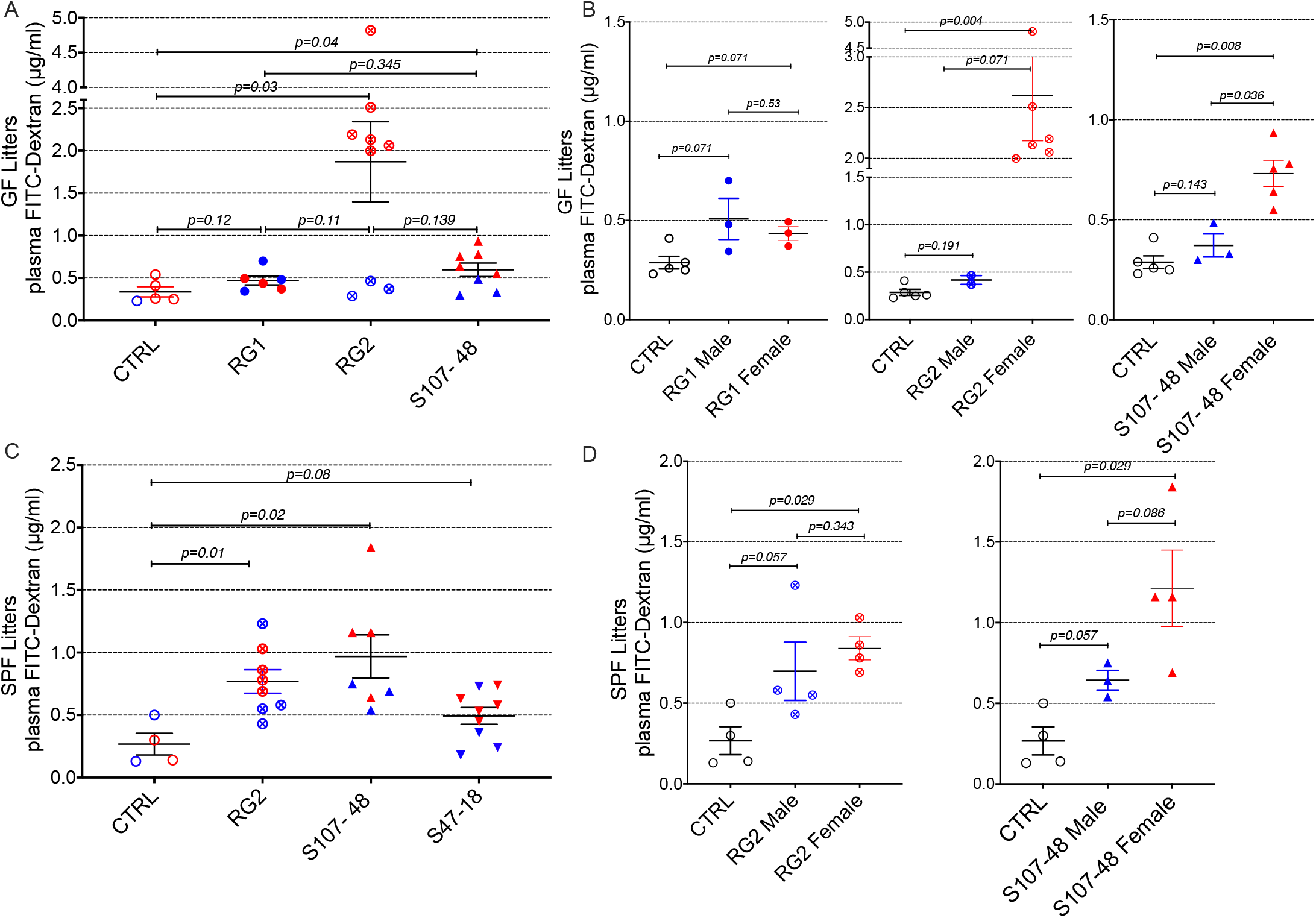
Intestinal colonization of GF and SPF mice by select RG strains induces enhanced intestinal permeability. A) Neonatal intestinal colonization of the litters of GF mice by RG strains, RG2 and S107-48 but not the RG1 strain, induces increased intestinal permeability when tested by the FITC-dextran challenge assay in mice at 10-14 weeks of age. B) Higher levels of intestinal permeability were found in RG2 and S107-48 strain colonized females than male littermates. C) Litters of SPF mice that were colonized by the RG strains, RG2 and S107-48, displayed increased intestinal permeability. Due to heterogeneity in responses the group colonized by the S47-48 RG strain did not attain significance. D) Female mice colonized with RG2 and S107-48 displayed significantly greater levels of intestinal permeability than male mice or non-colonized control mice. Results indicate mean ± SEM (n= 3-6 mice per group).

Mating of the pairs colonized with the different strains resulted in litters except for the RG1-colonized mice that did not yield progeny. In the litters of RG-colonized SPF mice, we again found evidence of increased gut permeability in the mice colonized by the RG2 and the Lupus S107-48 RG strains (Figure 3C). While there was a numerical trend toward greater intestinal permeability was found in the female progeny, compared to the male progeny, colonized by the RG2 strain. Compared to controls, a numerical trend towards higher mean levels was also documented in the male mice colonized by the Lupus S107-48 RG strain (*p*=0.057). Strikingly, significantly the highest mean levels of intestinal permeability were demonstrated in the female mice colonized with the Lupus S107-48 RG strain (*p*=0.029) (Figure 3D). These data demonstrate there is a female bias for induction of impaired gut barrier function resulting from colonization by the S107-48 RG Lupus strain.

### RG translocation to MLN in RG colonized mice with altered intestinal permeability

Based on above-described evidence of impaired gut barrier, we examined tissue extracts of colonized mice using the RG-specific 16S assay. Reiterating the patterns seen in the intestinal permeability assays (Figure 3), we found significant translocation of RG-specific genomic DNA in the female mice (*p*=0.001), with RG DNA levels also significantly above the levels found in male mice littermates colonized with the same S107-48 Lupus strain (*p*=0.002) (Figure 4). However, persistent RG DNA was not found in splenic extracts from the same mice (data not shown). The findings suggest that bacterial fragments can traverse the gut barrier as a consequence of colonization with some, but not other RG strains, and female mice are much more susceptible to RG-mediated breaches in the gut barrier.

**Figure 4.**
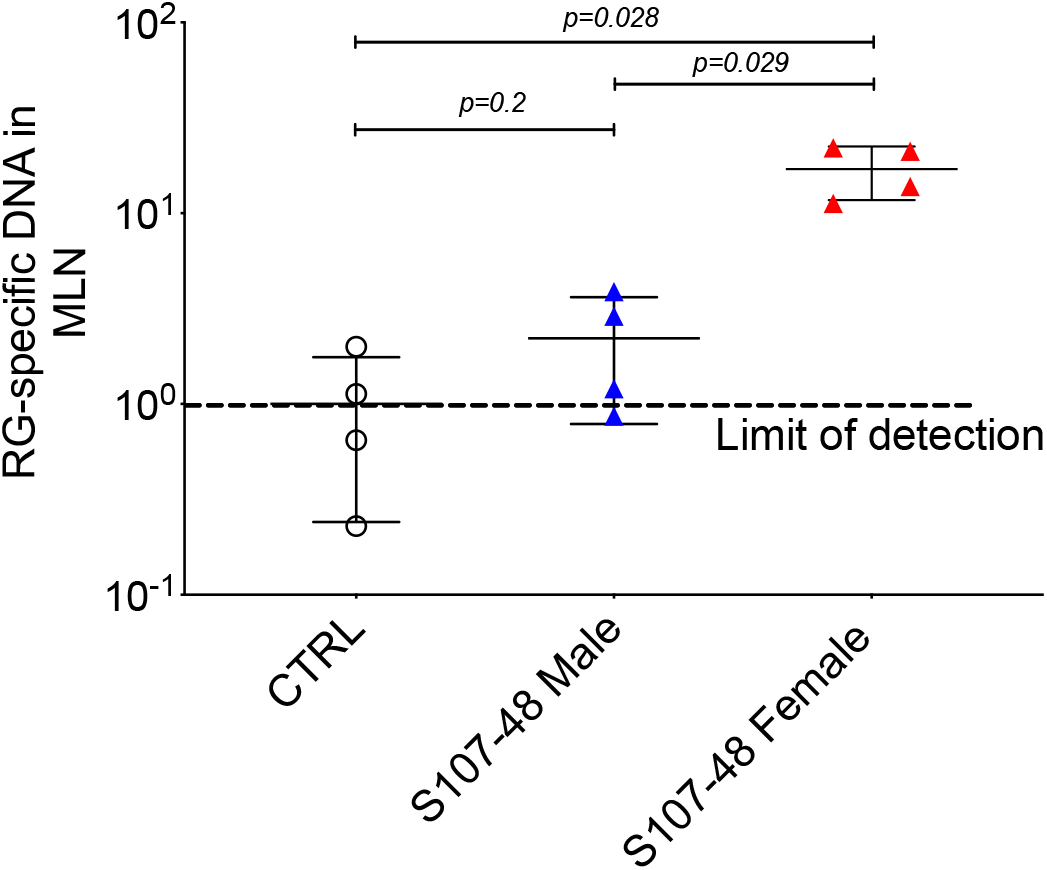
Translocation of S107-48 RG strain 16S rRNA gene DNA in female mice colonized by the S107-48 RG strain. Extracts of draining mesenteric lymph nodes (MLN) from female 14 wk-old littermates from GF mice colonized with the S107-48 RG strain demonstrate translocated RG genomic DNA (*p*=0.001). Levels in females were significantly higher than in males (*p*=0.002).

### Systemic Immune and autoimmune consequences of RG-mediated breaches of the gut barrier

Based on evidence of a breach in the gut barrier, we evaluated mice from litters of colonized germ-free mice for specific systemic immunorecognition of RG-associated antigens. In these studies, we found that RG colonization appeared to raise total IgG levels compared to noncolonized controls (Figure 5A), with significantly raised levels of serum IgG antibodies in S107-48 colonized mice that recognized the purified RG2 strain lipoglycan and the S47-18 strain lipoglycan antigens (Figure 5 B&C). In part, these findings are consistent with antigenic cross-reactivity between lipoglycans isolated from the RG2 and Lupus-associated RG strains (manuscript in preparation), while extracts of the RG1 strain were non-cross-reactive (1). Furthermore, there was no detectable reactivity with LPS isolated from Klebsiella or Salmonella (Figure 5 D&E), nor with pneumococcal cell wall C-polysaccharide (Figure 5F), which suggests that the anti-RG reactivity was antigen-specific.

**Figure 5.**
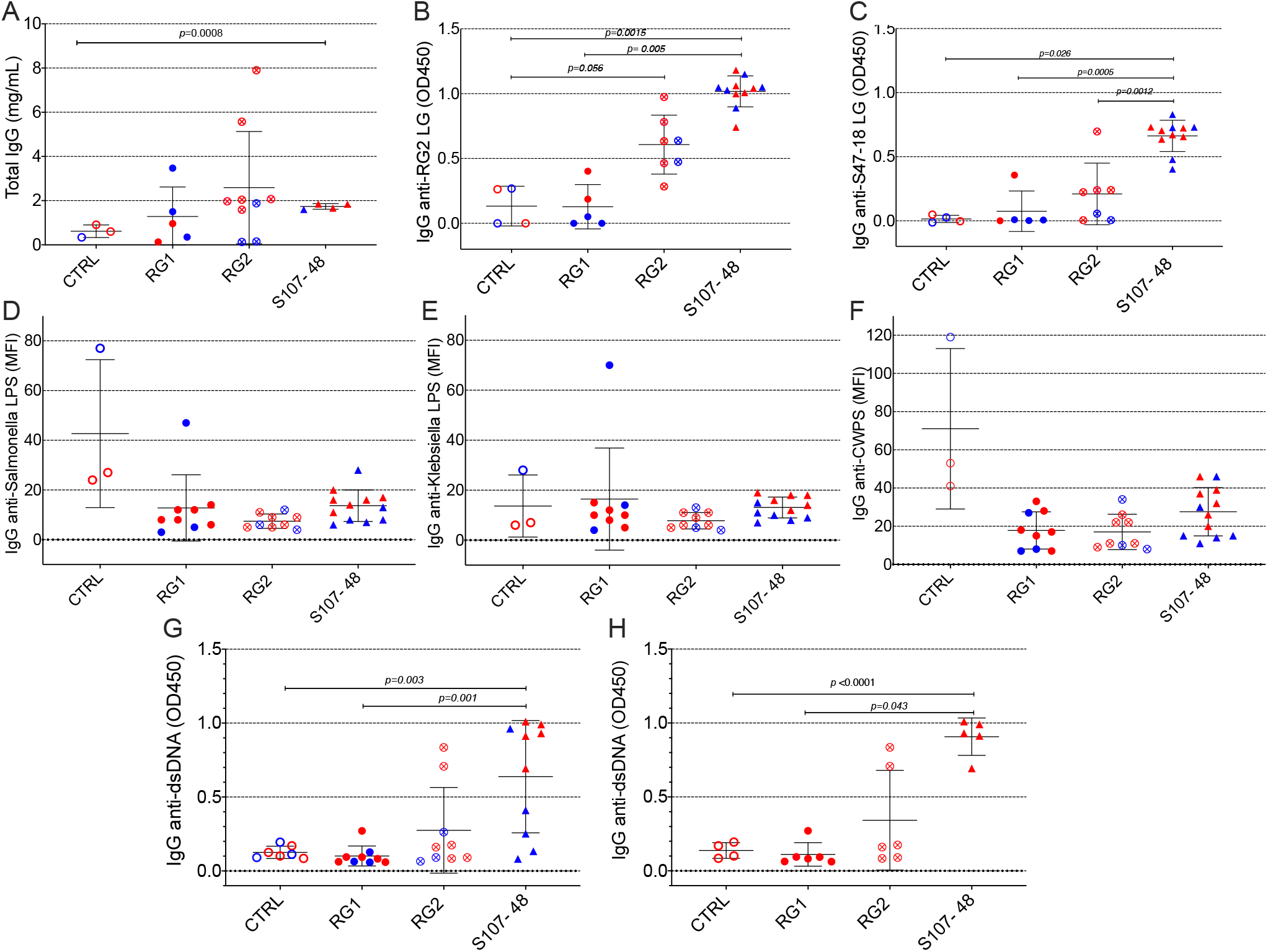
Intestinal colonization with certain RG strains induces IgG RG strain-associated cell wall lipoglycan antibodies and anti-native DNA autoantibodies. A) Following neonatal colonization of litters from RG colonized GF breeding pairs, S107-48 RG strain colonized mice display numerically higher mean serum total IgG levels, compared to controls. B) Neonatal colonization with the RG2 strain and the S107-48 RG strain induced significantly raised serum levels of IgG anti-RG2 strain cell wall lipoglycan antibodies. C) Neonatal RG colonization does **not** induce raised IgG-antibody levels to; D) Salmonella LPS, E) Klebsiella LPS, or F) to pneumococcal cell wall polysaccharide (CWPS). G) Neonatal colonization with the S107-48 RG strain induces raised serum levels of IgG anti-native DNA autoantibodies. H) Elevation of IgG anti native DNA following neonatal S107-48 RG strain was greatest in the female mice. (n= 4 to 11 per group). Antibody assays for IgG anti-native DNA used plasma diluted at 1:100 in ELISA (OD450). Antibody assays for IgG antibodies to RG lipoglycans, LPS and CWPS were performed by bead-based multiplex array (MFI) with plasma diluted 1:1000.

Based on the reported associations in Lupus patients between levels of circulating IgG anti-RG lipoglycan and Lupus anti-nuclear antibody production (ANA), we assayed plasma samples for binding reactivity with native thymic genomic DNA (dsDNA). These studies demonstrated significantly raised levels of IgG anti-DNA antibodies in mice after neonatal colonization with S107-48 Lupus strain (*p*=0.003)(Figure 5G), which was primarily due to the raised anti-DNA levels in the female mice (*p*<0.0001)(Figure 5H). Furthermore, in individual S107-48 colonized mice the level of intestinal permeability directly correlated with IgG anti-RG2 lipoglycan antibodies (Spearman r=0.7159, *p*=0.015) and with the level of IgG anti-native DNA autoantibodies ((r= 0.8679, *p*=0.0005)(Figure 6). These studies therefore document that female mice are more susceptible to Lupus RG induced impairment in the gut barrier, with immunologic consequences of induction of serum antibodies to RG strain-associated lipoglycan determinants, and for Lupus anti-DNA autoantibody production.

**Figure 6.**
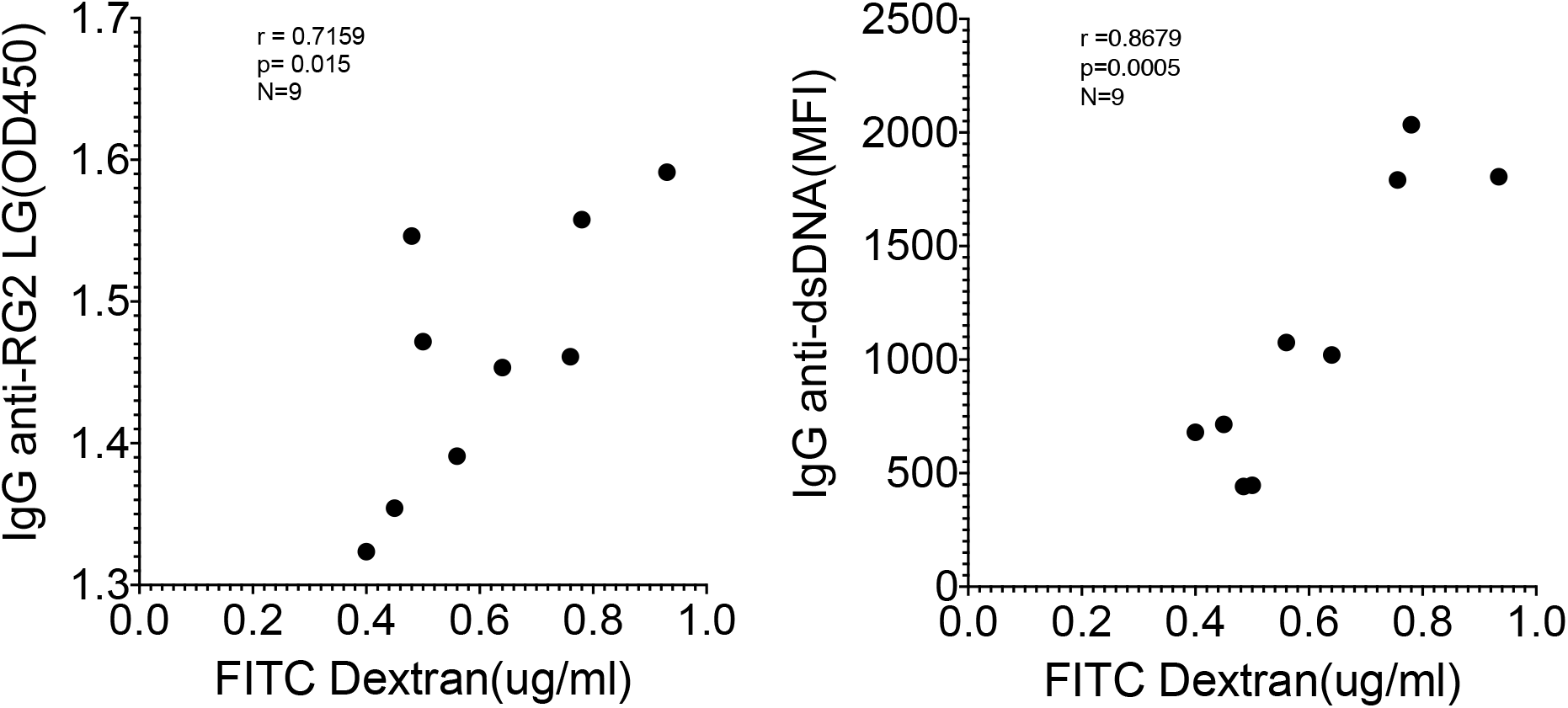
Correlation of levels of IgG anti-RG2 strain cell wall lipoglycan antibodies and IgG anti-native DNA autoantibodies correlate with levels of RG induced increased intestinal permeability. Each point represents an individual male and female littermate that had been neonatally colonized with S107-48 RG strain from a GF breeding pair, as depicted in Figures 5 B & G, respectively. Results from Spearman correlation analysis. IgG anti-RG2 plasma tested at 1:1000 dilution, and IgG anti-dsDNA was tested at a 1:100 dilution.

To further investigate the mechanisms responsible for increased gut permeability, at 14 weeks of age mice were sacrificed and sections of the small intestine were harvested and histopathologic examinations were performed. These preliminary studies demonstrated that unlike the expected morphology of intestinal villi and crypts seen in the control mice raised under GF conditions, that following neonatal colonization with the Lupus S107-48 RG strain the littermates of both sexes demonstrated histologic abnormalities with severely shortened epithelial villi and changes in the submucosa (Figure 7A). From RNA extracted from these small intestinal samples demonstrated trends towards raised levels of tight-junction Occludin gene transcripts in the RG2 colonized mice (*p*=0.06). Remarkably, there were significantly raised Occludin transcript levels in mice of both sexes colonized with the S107-48 Lupus strain (*p*=0.007) (Figure 7B). These findings are consistent with above-described evidence of functional intestinal barrier impairment induced by colonization with some RG strains.

**Figure 7.**
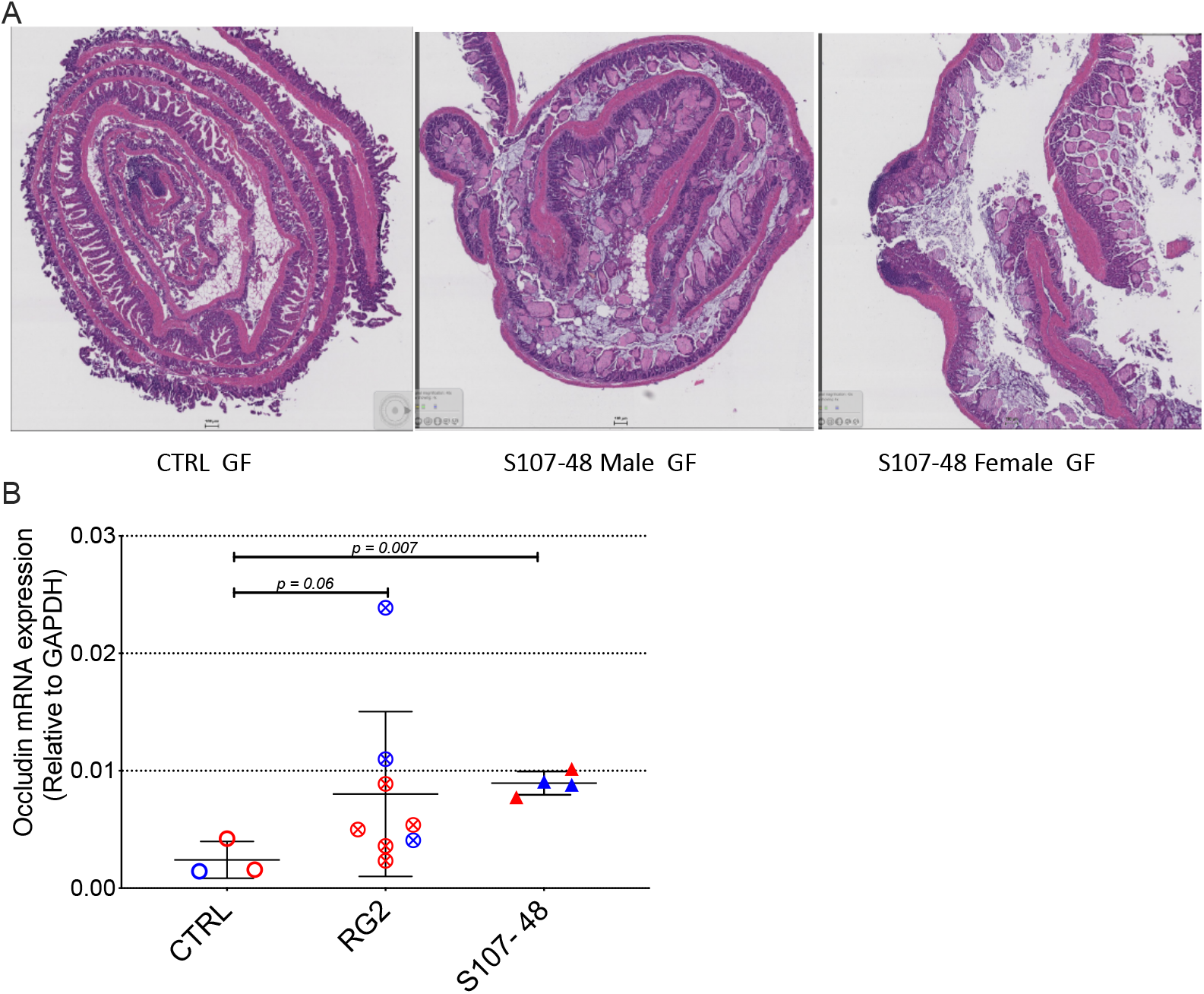
Neonatal RG colonization causes morphologic intestinal abnormalities and raised tight junction Occludin gene transcript levels. A) Representative H&E staining of terminal Ileum sections from litters of RG-colonized and control or GF mice, with comparisons of mice sacrificed at 14 weeks of age (n = 4 mice/group). B) Quantitative PCR of reverse transcribed (RT-qPCR) occludin gene transcripts from terminal ileum RNA extracts (n = 3 to 8 mice/group).

### RG strain-specific increases in gut permeability is reversible with oral treatment with larazotide

To next considered a potential causal role of the elevated serum zonulin levels in the functional abnormalities documented in S107-48 RG strain colonized mice, we repeated the FD4 challenge assay then treated groups of mice for ten days with oral administration of larazotide acetate (AT-1001), an octapeptide, originally identified from studies of the zonula occluden toxin by *Vibrio cholera* (24). Strikingly, larazotide treatment resulted in normalized gut barrier function, which became completely impervious to the passage of the fluorochrome-labeled dextran compound. Barrier function was normalized in both male and female mice, whether induced by the RG2 strain (*p*=0.03) or the Lupus S107-48 or Lupus S47-18 strains (*p*=0.01 for each)(Figure 8C&D), compared to control mice (Figure 8E). As larazotide is known to act locally to decrease tight junction (TJ) permeability by blocking zonulin receptors to promote TJ assembly and actin filament rearrangement (25), these findings document that gut colonization with Lupus RG strains, which results in intestinal permeability, with associated immune responses to microbial and nuclear antigens, can be specifically be inhibited by an agent with a known molecular pathway known to contribute to celiac disease and IBD (26).

**Figure 8.**
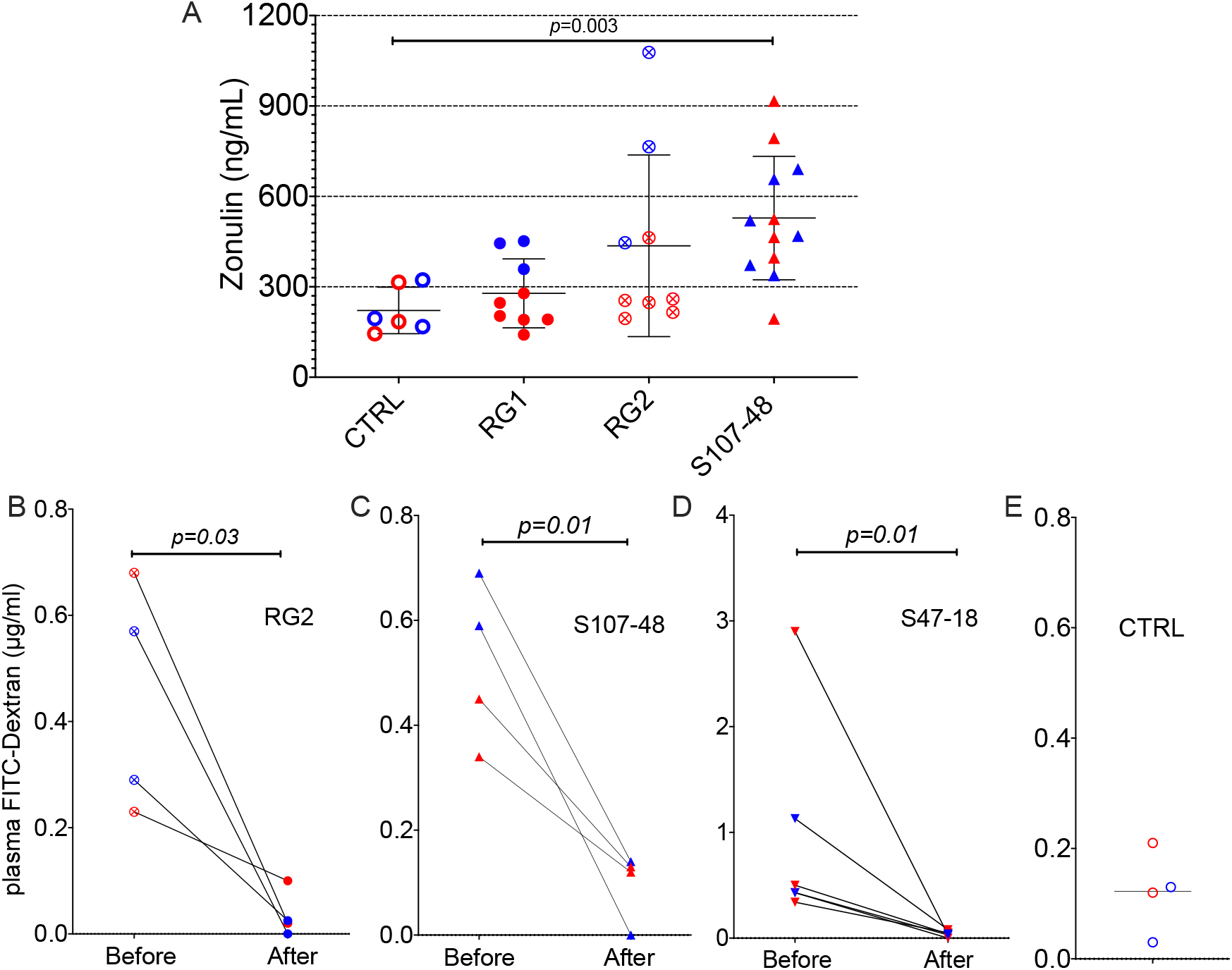
Oral larazotide treatment normalizes increased gut permeability induced by RG strain neonatal colonization. A) Plasma zonulin levels in 14-week-old mice after neonatal colonization, were significantly elevated in S107-48 RG strain mice compared to non-colonized controls (n = 5 to 12 mice). For each group mean+/-SD. In littermates from RG colonized SPF breeding pairs, individual mice were retested with results shown (before), then after a ten-day treatment with larazotide peptide in the water supply and 48 hr rest, gut permeability was then retested (after), with plasma FD4 levels shown for; B) RG2 strain colonized mice, C) S107-48 RG Lupus strain colonized mice, D) S47-18 RG Lupus strain colonized mice, with comparison to E) levels detected in non-colonized control mice. For the bottom panels, significance was tested by paired *t* test. *p<0.05 was considered statistically significant.

## Discussion

Whereas recent studies have documented that there are intestinal RG blooms in a subset of patients with active Lupus nephritis temporally associated with disease flares (1, 2), the current studies investigated the potential *in vivo* pathogenic properties of RG. As RG is common throughout healthy populations, albeit at lower levels, we reasoned that adverse local effects of intestinal RG colonization on the host may reflect quantitative expansions within intestinal communities and/or qualitative differences arising from genomic variations of these different RG strains. Importantly, diverse RG strains were able to colonize the murine GI tract, including after neonatal RG colonization that must have been mediated by the coprophagic behaviors of the litters from colonized breeding pairs. In part, because adult mice raised under GF conditions are known to have associated developmental immunodefects (reviewed in (27)) this finding was especially important as it enabled the study of mice that were neonatally colonized by a natural approach.

Our experimental design enabled comparisons of the influences of RG strains isolated from healthy donors, with strain(s) isolated from Lupus patients. To facilitate quantification of the level of RG colonization, we applied a previously reported RG species-specific genomic qPCR assay. This method was adopted due concern that different strains may differ in the efficiency of in vitro culture, which could affect quantitation of recovered colony forming units (CFU), albeit we adopted a method that cannot discriminate between viable and no longer viable bacterial cells. In any case, our methodologic approach allowed us to address an enigma highlighted by studies of murine colonization by *Enterococcus gallinarum,* which found small intestinal colonization in the absence of detectable *E. gallinarum* in the fecal pellets (28). In our studies, individual mice with persistent and substantial level of RG colonization, based on RG detection in cecal luminal contents, at times had no detectable levels RG in matched fecal samples. These findings provide a nuanced technical insight into the limits of 16S rRNA gene amplicon analysis of fecal samples which has become a widely accepted and unbiased approach to gut microbiota community analysis.

Our studies demonstrated that colonization with some RG strains, which included CC55_001C (RG2) as well as the S107-48 and the S47-18 Lupus strains reproducibly induced enhanced gut permeability. Unexpectedly we found that despite effective colonization the type-specific RG strain, VC1-7 (RG1) did not appear to affect the gut barrier. Arguably, the benign influence of this strain could be linked to the reported anti-inflammatory properties of the capsular polysaccharide expressed by this strain from a healthy donor (11). While the Lupus strains, and also RG2 strain each induced increased intestinal permeability, a specific molecular mediator for these host functional effects was not elucidated and will be the focus of future investigations.

Although there was no consistent pattern of sex-biased intestinal colonization, female mice consistently demonstrated higher levels of intestinal permeability, whether as a consequence of colonization of GF mice, neonatal colonization or of antibiotic-conditioned mice raised under SPF conditions. These findings may be especially relevant to the role of RG blooms during Lupus pathogenesis, a condition that affects nine-fold more women than men. Notably, recent reports have shown that a major subset of female Lupus patients have subclinical abnormalities attributed to breaches in the gut barrier (1) as well as their first degree female relatives (29), and we wonder whether the same mechanisms are responsible in patients as we found in our murine colonization studies.

Amongst the most significant finding was that impairments of the intestinal barrier defect induced by Lupus RG strains was associated with raised serum levels of zonulin, the only known physiological regulator of intestinal intracellular tight junctions (24). While first discovered in studies of gluten enteropathy, intestinal bacteria (including both pathogens and certain commensals) have also been identified as stimuli that can trigger the release of zonulin (30). Strikingly, the abnormal results of the FITC-dextran assays were completely normalized by oral administration of the specific zonulin receptor antagonist, the octapeptide larazotide was originally identified from studies of the zonula occludens toxin (ZOT) secreted by *Vibrio cholera* (25).

In our studies, post-challenge levels of FD4 that measured intestinal permeability directly correlated with the levels of IgG antibodies to cell wall lipoglycan purified from an RG strain that increased murine gut permeability (Figure 6). There was also induction of IgG antibody native DNA antibodies that are a diagnostic criterion for clinical SLE (31). The current findings document another clinically relevant model in which zonulin plays an important role in the balance between tolerance and immunity (26), and our studies also provide mechanistic rationalization for findings in Lupus nephritis patients that fecal RG abundance correlated with levels of serum IgG antibodies to RG2 lipoglycan (1). Taken together, our findings suggest a molecular mechanism by which strains of RG that colonize Lupus patients may be responsible for inducing defects in the gut barrier that contribute to autoantibody and immune complex formation that are known to be key drivers in the pathogenesis of Lupus Nephritis (32). Moreover, larazotide has been shown to be safe in early trials, and is now being evaluated in two Phase III trials in celiac disease (33, 34). We therefore propose that larazotide should also be considered as a potential adjunct therapeutic approach for blocking the harmful influence of gut dysbiosis during Lupus immunopathogenesis.

---

**Supplementary Figure 1.**
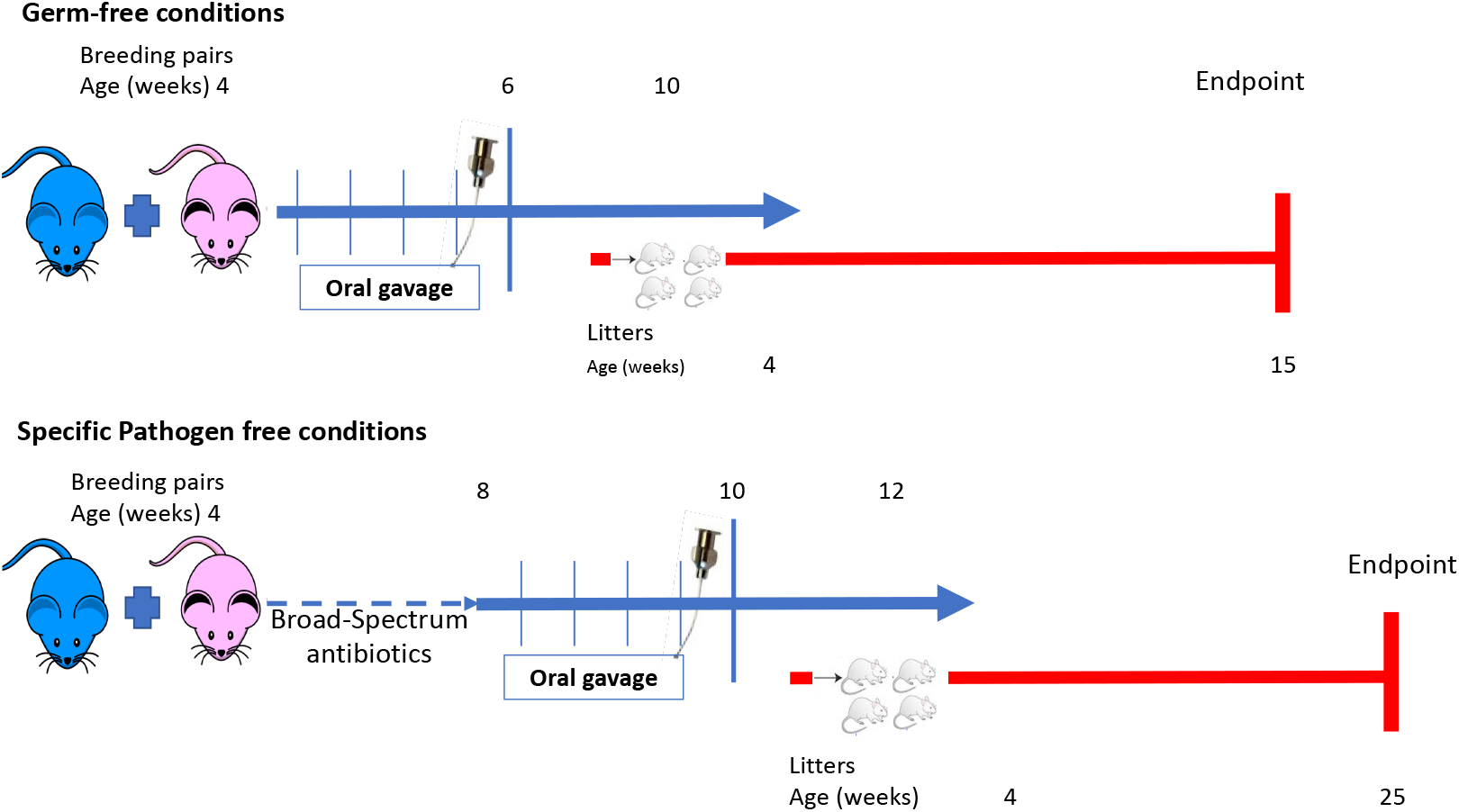
Overview of murine RG colonization studies.

## Acknowledgments

We acknowledge the assistance of the NYU Histology Core, the NYU Microscopy Laboratory and the NYU Genome Technology Core.

